# Decreased calmodulin recruitment triggers PMCA4 dysfunction and pancreatic injury in cystic fibrosis

**DOI:** 10.1101/2020.09.10.290940

**Authors:** Tamara Madácsy, Árpad Varga, Noémi Papp, Barnabás Deák, Bálint Tél, Petra Pallagi, Viktória Szabó, Júlia Fanczal, Zoltan Rakonczay, Zsolt Rázga, Meike Hohwieler, Alexander Kleger, Mike Gray, Péter Hegyi, József Maléth

## Abstract

Exocrine pancreatic damage is a common complication of cystic fibrosis (CF), which can significantly debilitate the quality of life and life expectancy of CF patients. The cystic fibrosis transmembrane conductance regulator (CFTR) has a major role in pancreatic ductal ion secretion, however, it presumably has an influence on intracellular signaling as well. Here we describe in multiple model systems, including iPSC-derived human pancreatic organoids from CF patients, that the activity of PMCA4 is impaired by the decreased expression of CFTR in ductal cells. The regulation of PMCA4, which colocalizes and physically interacts with CFTR on the apical membrane of the ductal cells, is dependent on the calmodulin binding ability of CFTR. Moreover, CFTR seems to be involved in the process of the apical recruitment of calmodulin, which enhances its role in calcium signaling and homeostasis. Sustained intracellular Ca^2+^ elevation in CFTR KO cells undermined the mitochondrial function and increased apoptosis. Based on these, the prevention of sustained intracellular Ca^2+^ overload may improve the exocrine pancreatic function and may have a potential therapeutic aspect in CF.

## Introduction

Cystic fibrosis (CF) is the most common autosomal recessive genetic disease characterized by multiorgan pathology due to impaired function of all cystic fibrosis transmembrane conductance regulator (CFTR) expressing epithelia^1,2^. Life expectancy of CF patients has increased due to recent therapeutic advances, however the average age of death still remained 31.4 years^3^. With the increasing disease duration the gastrointestinal manifestations of CF, which include exocrine pancreatic insufficiency (EPI) in the majority of cases (83%)^3^ and CF-related acute pancreatitis (AP) with a consequent malnutrition come into prominence^4^. The destruction of the gland starts *in utero* due to the secretory defect resulting in the production of viscous, protein rich fluid leading to protein precipitation and ductal obstruction by mucoprotein plugs, inflammation, acinar destruction, cyst formation and eventually generalized fibrosis and calcification^2^. Importantly, ~20 % of the patients have evidence of pancreatic damage but retain sufficient exocrine pancreatic function to maintain digestion (exocrine pancreatic sufficient - EPS), however EPS patients are at an increased risk of developing recurrent AP^2,5^. The damage of the exocrine pancreas will lead to the destruction of the islets of Langerhans promoting to the development of CF-related diabetes^6^, which associates with increased mortality in CF population^7^ In addition, a recent study by Sun et al. suggests that in utero and postnatal administration of VX-770 (a CFTR channel potentiator) partially prevents the development of multiorgan damage, including pancreatic injury, in CFTR knockout ferrets^8^. These results suggest that the percentage of the EPS patients may increase soon.

CFTR is expressed on the apical membrane of the pancreatic ductal epithelial cells in the exocrine pancreas and plays a fundamental role in the secretion of alkaline pancreatic juice^9^. The physiological significance of the exocrine pancreatic secretion is well-established by independent investigations^9,10^ The HCO_3_^-^-rich fluid washes out digestive proenzymes from the ductal lumen and neutralises the protons co-released during exocytosis by acinar cells^11^. Moreover, the alkaline intraductal pH is crucial to prevent the premature intrapancreatic trypsinogen autoactivation^12^. Dysfunction of the ductal secretion leads to disturbed plasma membrane dynamics in the pancreatic acinar cells of CFTR knockout mice^13^. In addition, the role of CFTR in the development of ethanol-induced AP, which is one of the most common form of AP, was demonstrated by our group. We provided direct evidence that ethanol and fatty acids reduce the activity and expression of CFTR in pancreatic ductal cells leading to impaired HCO3^-^ secretion, which increased the severity of alcohol-induced AP^14^. Importantly, pharmacological restoration of CFTR mediated secretion with the corrector C18 and the potentiator VX-770 restored acinar cell function and decreased pancreatic inflammation in mouse models of chronic and autoimmune pancreatitis^15^. As demonstrated by these studies, the secretory function of CFTR is crucial to maintain the proper function of the exocrine pancreas. However, CFTR has also been suggested by recent studies to possess a more complex integrative role in the regulation of intracellular signalling events in addition to the secretion of Cl^-^ and HCO3^-^ ions, which is much less characterized in epithelial cells. In cultured airway epithelial cells increased IP3R-dependent Ca^2+^ release^16^, elevated activity of the sarco/endoplasmic reticulum Ca^2+^-ATPase (SERCA) pump and enhanced mitochondrial Ca^2+^ uptake were described in CF, whereas plasma membrane Ca^2+^ ATPase (PMCA) function was decreased^17^ However, the exact mechanism of these changes is not established yet. Notably, similar alterations of the intracellular Ca^2+^ signalling (such as sustained intracellular Ca^2+^ overload, impaired SERCA and PMCA functions) were observed in the pancreatic acinar and ductal cells during AP^18^, which raises the possibility that similar changes can have a significant impact on the exocrine pancreas in CF. However, currently we have no information how the intracellular signalling changes in the ductal epithelia and how this affects ductal function in CF.

Therefore, in this study we investigated the Ca^2+^ homeostasis of CFTR KO pancreatic ductal cells using multiple independent model systems, including iPSC-derived human pancreatic organoids. We demonstrated that the activity of PMCA4 is impaired in the lack of CFTR expression resulting in sustained elevation of the intracellular Ca^2+^ concentration. We also demonstrated that PMCA4 colocalizes and physically interacts with CFTR on the apical membrane of the ductal cells. The regulation of PMCA4 activity seems to involve recruitment of calmodulin to the apical membrane by CFTR. Finally, we demonstrated that the sustained elevation of intracellular Ca^2+^ in CF ductal cells inhibit the mitochondrial function accompanied by an increase in apoptosis. Thus, our results suggest that the prevention of sustained intracellular Ca^2+^ overload may improve the exocrine pancreatic function in CF-related pancreatic damage.

## Results

### The function of plasma membrane Ca^2+^ pump is impaired in the absence of CFTR in pancreatic ductal cells

First, we wanted to compare the intracellular Ca^2+^ signalling in pancreatic ductal cells isolated from WT and CFTR KO mice. Therefore, to release the endoplasmic reticulum (ER) Ca^2+^ stores and trigger extracellular Ca^2+^ entry, we treated isolated ducts with 100 μM carbachol, which evoked a two-phase intracellular Ca^2+^ elevation both in WT and CFTR KO ductal cells (Figure 1. A.). The maximal Ca^2+^ release was not different, however we found that the slope of the plateau phase – representing the Ca^2+^ extrusion from the cytosol - was significantly increased in CFTR KO vs WT ducts. To rule out that the observed difference is due to nonspecific tissue damage in CFTR KO mice we used isolated acinar cells, which do not express CFTR. We did not see any difference in the maximal intracellular Ca^2+^ elevation, or in the slope of plateau phase upon carbachol treatment between acinar clusters of WT and CFTR KO mice (Figure 1. B.) revealing that the alteration in ductal cells is due to the decreased CFTR function, or expression. To clarify whether the impaired of CFTR function or expression caused the alteration in the intracellular Ca^2+^ signalling we used a selective pharmacological inhibitor of CFTR. 10 μM CFTR(inh)-172 significantly decreased the intracellular pH regeneration of isolated ducts after alkali load (Supplementary Figure 1. A.) suggesting the inhibition of CFTR function^14^. Notably, the carbachol-induced intracellular Ca^2+^ signalling was not different in 10 μM CFTR(inh)-172 treated WT ductal cells (Figure 1. C.) suggesting that the lack of CFTR function has no effect on the intracellular Ca^2+^ signalling. Next, we induced complete ER Ca^2+^ store depletion with 25μM cyclopoazonic-acid (CPA) in Ca^2+^-free extracellular media. Under these conditions the store depletion activates the store operated Ca^2+^ influx (SOCE) and the re-addition of extracellular Ca^2+^ induces a SOCE-mediated rise in the intracellular Ca^2+^ concentration. The inhibition of SERCA pump with CPA caused rapid release of the ER Ca^2+^ stores, which was not different in WT and CFTR KO cells (Figure 1. D). To evaluate Ca^2+^ extrusion we removed the extracellular Ca^2+^ again, which resulted in a rapid drop of intracellular Ca^2+^ concentration. Under these conditions, the slope of Ca^2+^ efflux was significantly decreased in CFTR KO ductal cells. To avoid any potential errors in the evaluation, the slope of the Ca^2+^ extrusion was compared at the same intracellular Ca^2+^ levels. As Ca^2+^ extrusion in non-excitable cells can be mediated by the activity of PMCA and Na^+^/Ca^2+^ exchanger (NCX), we used two inhibitors to rule out the contribution of NCX to Ca^2+^ extrusion. Neither the pan-NCX inhibitor CB-DMB nor SEA0400 had any effect (Supplementary Figure 1. B.), suggesting that NCX activity has no contribution to the Ca^2+^ efflux in ductal cells, and thus the activity of PMCA is impaired in CFTR KO ductal epithelial cells.

**Figure 1.**
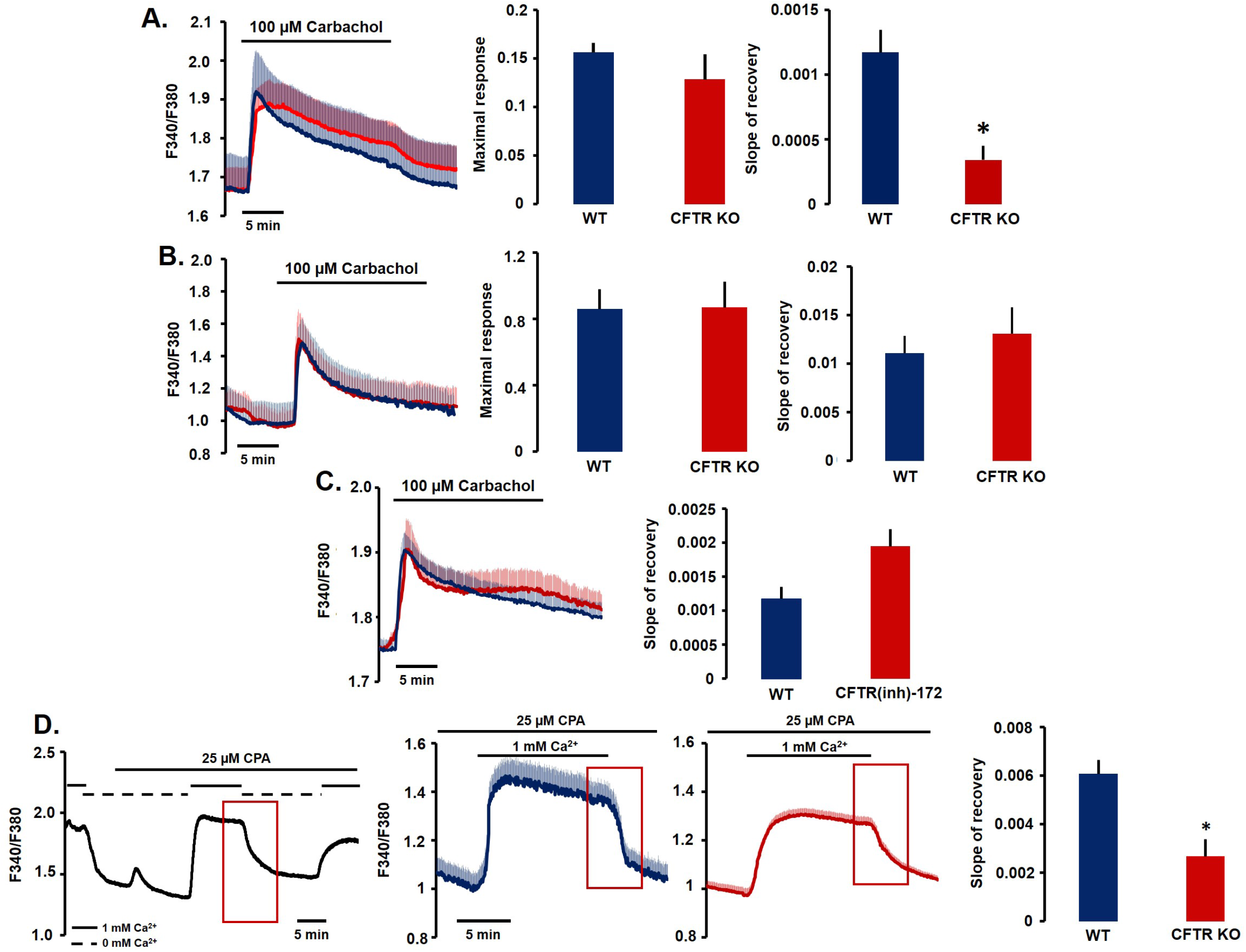
PMCA dysfunction in CFTR knockout pancreatic ductal cells. **A.** Average traces, maximal Ca^2+^ elevation and the slope of recovery in WT and CFTR KO pancreatic ductal fragments in response to 100 μM carbachol. The maximal Ca^2+^ release was not different, however Ca^2+^ extrusion was significantly impaired in CFTR KO ducts. **B.** Average traces, maximal Ca^2+^ elevation and the slope of recovery in WT and CFTR KO pancreatic acinar cells in response to 100 μM carbachol showed no difference. **C.** Average traces and the slope of recovery demonstrates that inhibition of CFTR with 10 μM CFTR(inh)-172 has no effect on the response to carbachol in WT pancreatic ductal fragments. **D.** Complete ER Ca^2+^ store depletion was induced with 25 μM cyclopoazonic-acid (CPA) in Ca^2+^-free extracellular media. Re-addition of extracellular Ca^2+^ induced a SOCE-mediated rise in the intracellular Ca^2+^, whereas Ca^2+^ extracellular removal trigger Ca^2+^ extrusion from the cytosol (red quadrant highlights the Ca^2+^ extrusion, which was used to calculate the slope of recovery). Under these conditions, the slope of Ca^2+^ efflux was significantly decreased in CFTR KO vs WT ductal cells. All averages were calculated from 6-10 individual experiments. *: p< 0.05 vs WT.

### CFTR knockdown impairs, whereas CFTR overexpression increases PMCA function

Next, we challenged our hypothesis further in the human pancreatic ductal cell line CFPAC-1, which was derived from the liver metastasis of a ductal adenocarcinoma from a patient with CF. We compared the PMCA function of CFPAC-1 cells with CFPAC-1 cells in which the CFTR expression has been corrected with Sendai virus mediated gene delivery of the human CFTR gene^19^. The restoration of the CFTR protein expression was confirmed by immunolabeling of CFTR (Figure 2. A.). We found that the correction of CFTR expression also restored PMCA function (Figure 2. B.). Next, we knocked down the CFTR expression in ductal fragments isolated from mouse with siRNA specific to CFTR. The protein expression markedly decreased (Figure 2. C.) and in the siCFTR treated ductal fragments (compared to siGLO Green treated controls) we detected impaired Ca^2+^ extrusion due to the diminished PMCA function (Figure 2. D.).

**Figure 2.**
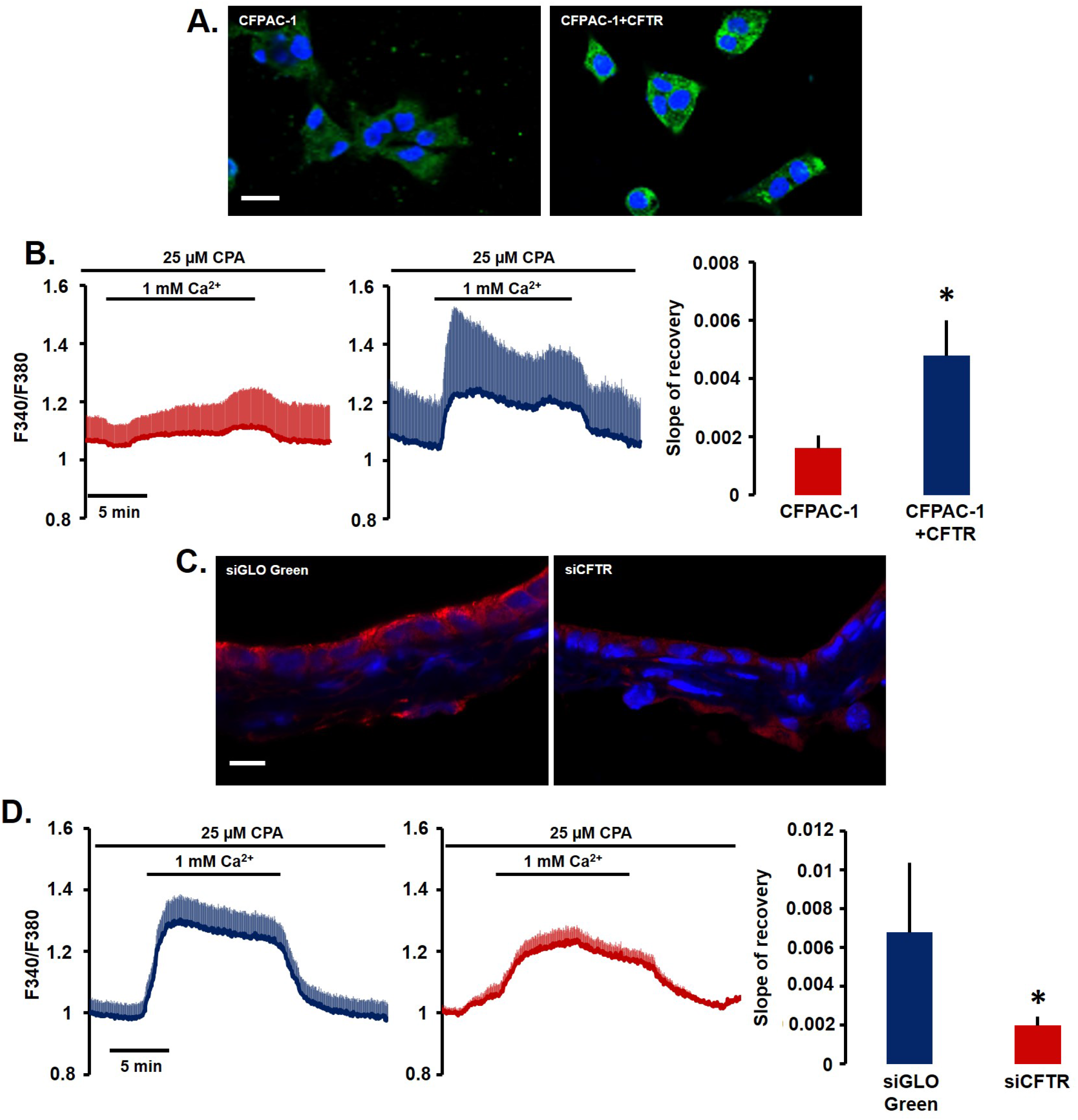
The effect CFTR overexpression and knockdown on the PMCA function in pancreatic ductal cells. **A.** Restoration of CFTR expression in CFPAC-1 human pancreatic ductal cell line CFPAC-1 with Sendai virus mediated gene delivery was confirmed by immunostaining. **B.** Average traces of intracellular Ca^2+^ concentration and bar chart of the slope of recovery highlight that the correction of CFTR expression also improved PMCA function. **C.** siRNA-mediated knockdown in isolated mouse ductal fragments markedly decreased CFTR expression. **D.** Average traces of intracellular Ca^2+^ concentration and bar chart of the slope of recovery demonstrate that the siCFTR treatment significantly decreased the PMCA function. Scale bars: 10 μm. All averages were calculated from 6-10 individual experiments. *: p< 0.05 vs CFPAC-1, or siGLO Green, respectively.

### PMCA4 colocalizes with CFTR at the apical membrane of pancreatic ductal cells

Reverse transcription polymerase chain reaction (RT-PCR) followed by endpoint analysis revealed strong expression of PMCA 1 and 4 both in whole pancreatic tissue and isolated mouse pancreatic ducts (Figure 3. A.). The PMCA2 gene showed low, but detectable expression in both samples. To compare the relative expression levels of PMCA genes we performed quantitative real-time PCR (qRT-PCR) in the isolated ductal fragments of WT and CFTR KO mice (Figure 3. B.), which showed no significant difference of PMCA1 and 4 expression. Next, we determined the expression of PMCA1 and 4 proteins in isolated pancreatic ducts with immunofluorescent labelling (Figure 3. C.). According to our results PMCA4 is expressed on the apical membrane of the pancreatic ductal epithelial cell, whereas PMCA1 showed an even distribution in the apical and basolateral membrane. The expression of PMCA1 and 4 proteins was not altered in CFTR KO ducts, whereas a strong colocalization of PMCA4 and CFTR was visible at the apical membrane with a Mander’s overlap coefficient of 0.906 (Figure 3. D.). These results suggest that the lack of CFTR expression diminishes the function, but not the expression of PMCA4 in ductal epithelial cells.

**Figure 3.**
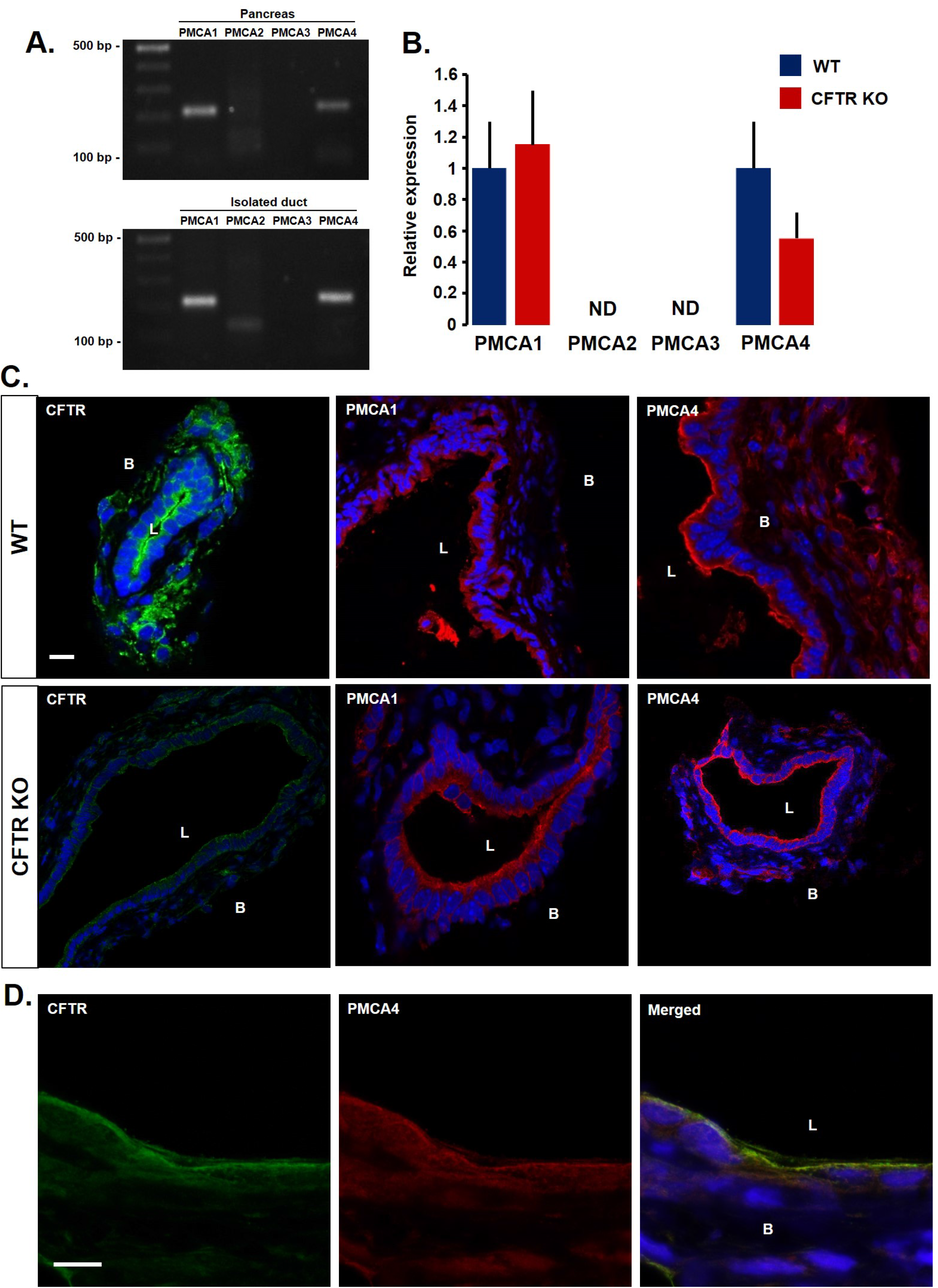
The expression of PMCA isoforms in mouse pancreatic ductal cells. **A.** Endpoint RT-PCR analysis of PMCA isoforms in whole pancreas (upper panel) and in isolated ducts (lower panel). The expression of PMCA1 and 4 was detected. **B.** PMCA1 and 4 expression showed no significant difference in WT and CFTR KO pancreatic ductal fragments, whereas PMCA2 and 3 could not be detected (ND). **C.** The expression of PMCA1 and 4 in isolated pancreatic ducts. PMCA4 is expressed on the apical membrane of the pancreatic ductal epithelia, whereas PMCA1 showed an even distribution in the plasma membrane. **D.** PMCA4 and CFTR colocalize at the apical membrane with a Mander’s overlap coefficient of 0.906. Scale bars: 10 μm. B: basolateral side; L: lumen.

### Human CF pancreatic iPSC organoids recapitulate the alteration of PMCA function

To assess the human relevance and rule out the potential species specificity of our findings, we utilized human induced pluripotent stem cell (iPSC) derived pancreatic organoids^20^. CF-iPS cell lines from a donors affected by classical CF were established by lentiviral reprogramming of patient keratinocytes as previously published. Stepwise *in vitro* differentiation was performed to direct the iPSCs towards the pancreatic lineage and generate exocrine pancreatic organoids in 3D-suspension culture (Figure 4. A.). First, we analysed the expression of CFTR and PMCA4 in organoids established from control and CF patients (Figure 4. B.). Like isolated mouse ductal fragments both CFTR and PMCA4 were expressed in the control sample, whereas the CF organoids showed no staining for CFTR, which was markedly restored by 12 h incubation with the CFTR corrector VX-809 (10 μM). Importantly, the expression of PMCA4 was not changed in CF organoids. Next, we performed Ca^2+^ removal after ER depletion as described above and found a significant decrease in the Ca^2+^ extrusion in CF organoids compared to control, which perfectly recapitulates our observation in other model systems (Figure 4. C.). Importantly, pre-treatment of CF organoids with 10 μM VX-809 for 12 h significantly improved the Ca^2+^ extrusion also suggesting that in human CF organoids the decrease of PMCA function can be restored with CFTR corrector treatment.

**Figure 4.**
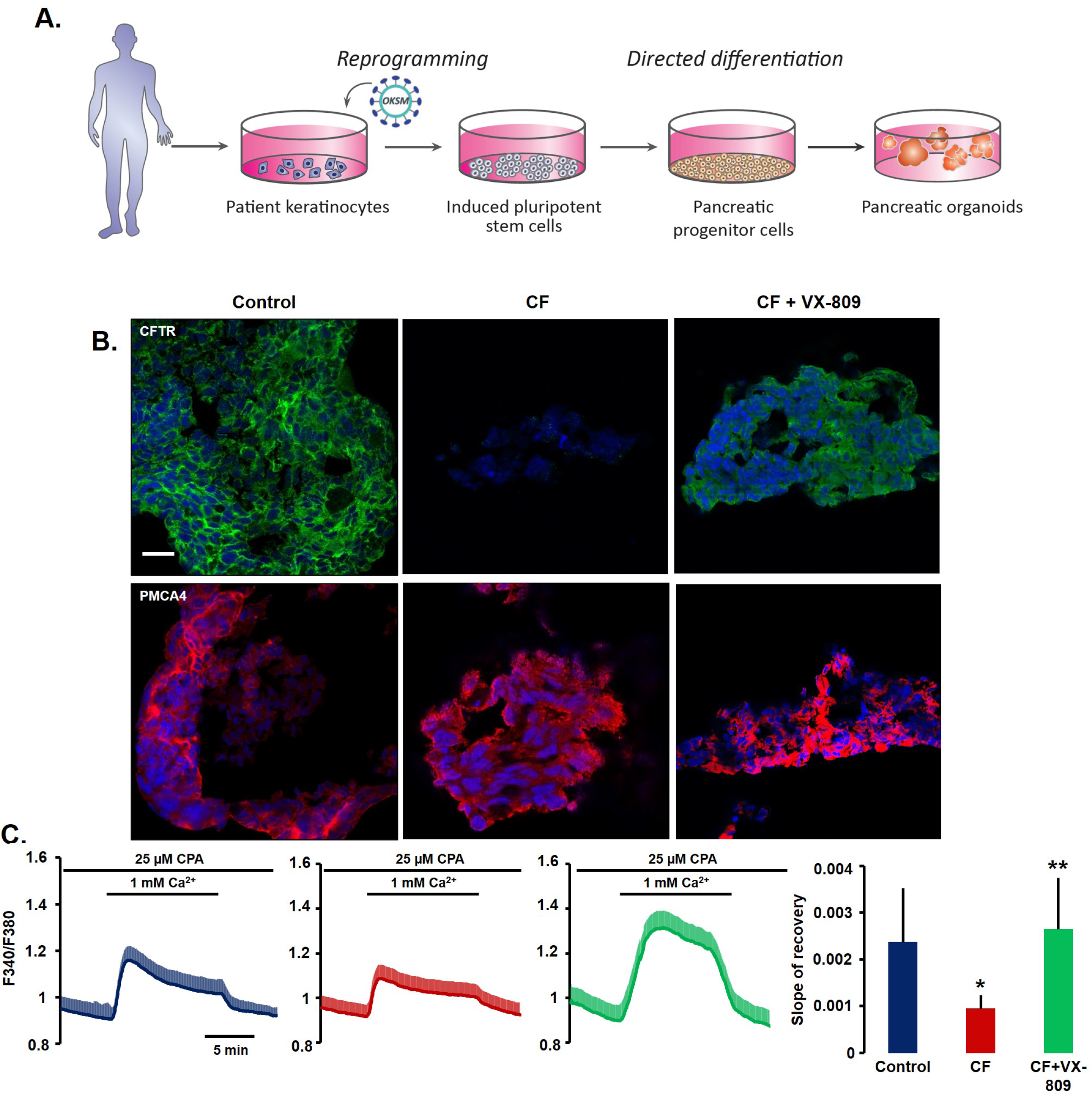
PMCA function in iPSC-derived pancreatic organoids from healthy controls and CF patients. **A.** Schematic outline of the generation of iPSC-derived pancreatic organoids. **B.** CFTR and PMCA4 were expressed in the control sample, whereas the CF organoids showed no staining for CFTR, whereas PMCA4 expression was not changed. 12 h incubation with 10 μM VX-809 restored the CFTR expression in the CF samples. **C.** Average traces and bar chart of the slope of recovery highlights a significant decrease in the Ca^2+^ extrusion in CF organoids compared to control, which was markedly improved by the CFTR corrector treatment. Scale bars: 10 μm. All averages were calculated from 6-10 individual experiments. *: p< 0.05 vs Control, **: p< 0.05 vs CF.

### PMCA4 interacts with CFTR at the apical membrane of pancreatic ductal epithelial cells

Our measurements suggested a close connection between CFTR and PMCA4, which seems to be required for the proper function of PMCA4 and for physiological Ca^2+^ extrusion. To characterize this connection further in cellular context Duolink proximity ligation assay (PLA) of endogenous PMCA and CFTR was performed. To avoid unspecific antibody binding, we used pancreatic ductal fragments isolated from guinea pig in this experiment. The expressions of PMCA4 and CFTR in guinea pig recapitulated the expression pattern of mouse ductal cells (Supplementary Figure 2.). The Duolink PLA probe suggested that PMCA4 and CFTR are in a proximity of <40 nm (Figure 5. A.). To be able to visualize this interaction at even higher resolution, we utilized the dSTORM technique first in HeLa cells cotransfected with plasmids encoding CFTR and PMCA4. This technique revealed a perfect overlap (<20 nm) between the two proteins in the plasma membrane suggesting a physical interaction between them (Figure 5.B., Supplementary Figure 3.). Next, we used 2D adherent primary human ductal cells generated from human pancreatic ductal organoids to confirm this interaction in endogenously expressed protein as well (Figure 5. C.). Similar to HeLa cells, the overlap of CFTR and PMCA4 was also confirmed in primary human pancreatic ductal cells (Figure 5. D.).

**Figure 5.**
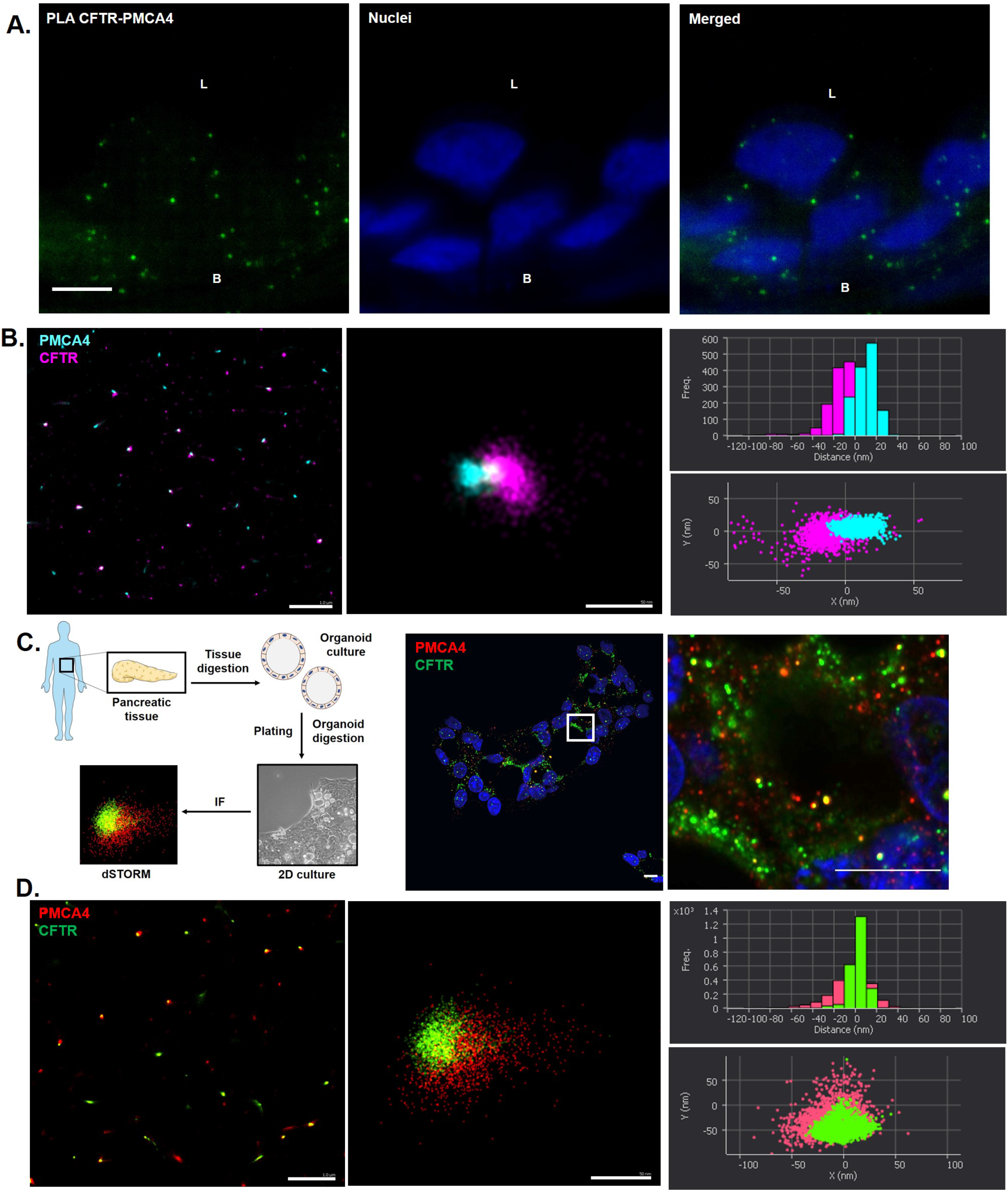
Intimate interaction of PMCA4 with CFTR in the plasma membrane. **A.** Duolink proximity ligation assay (PLA) of endogenous PMCA and CFTR was performed to assess the interaction of the two proteins. For representation, 18 images of a Z-stack were merged. The PLA suggested that PMCA4 and CFTR are in a proximity of <40 nm. Scale bars: 10 μm. **B.** dSTORM images of HeLa cells transfected with CFTR and PMCA4. This technique revealed a perfect overlap (<20 nm) between the two proteins in the plasma membrane. Scale bars: 1 μm and 50 nm, respectively. **C.** Confocal images of 2D adherent primary ductal cells generated from human pancreatic ductal organoids. Scale bars: 5 μm. **D.** dSTORM images of CFTR and PMCA4 in primary human pancreatic ductal cells demonstrating a strong colocalization of native proteins. Scale bars: 1 μm and 50 nm respectively. n= 5-7 cells were analysed for each condition in dSTORM.

### Calmodulin binding by CFTR regulates PMCA4 activity in pancreatic ductal cells

In the next step we wanted to provide more details about the connection of CFTR and PMCA4. A recent study by Bozoky et al. showed that calmodulin interacts with CFTR in the alternative binding conformation, in which the two lobes of calmodulin can bind to two separate sequences independently and bridge over larger distances^21^. Thus, this mode of calmodulin binding to CFTR might determine the activity of other calmodulin-regulated proteins, such as PMCA4. Indeed, calmodulin showed a strong colocalization with CFTR and PMCA4 at the apical membrane of ductal epithelial cells when investigated with dSTORM on cross-sections of mouse pancreatic ductal organoids (Figure 6. A.). To compare the intracellular distribution of calmodulin in WT and CFTR KO ductal cells, we stained calmodulin in cross-sections of isolated ductal fragments. Interestingly, calmodulin was strongly associated with the apical membrane in ductal epithelial cells of WT ductal fragments (Figure 6. B.). In contrast, in CFTR KO cells calmodulin was distributed evenly at the apical pole without association to the apical membrane as suggested by the line intensity profiles. Finally, we wanted to provide evidence that the lack of calmodulin binding by CFTR can impair the function of PMCA4 in CF. General knockdown, or inhibition of calmodulin can have multiple effects in the cells, therefore we cotransfected HEK-293 cells with WT or calmodulin binding site mutant CFTR(S768A) and PMCA4. Both WT and mutant CFTR were localised to the plasma membrane and colocalized with calmodulin (Figure 6. C.). Cotransfection of PMCA4 and CFTR markedly increased the slope of Ca^2+^ extrusion, whereas PMCA4 alone showed moderate activity (Figure 6. D.). More importantly, when compared to the WT CFTR, in cells transfected with CFTR(S768A) the PMCA4 activity was significantly impaired suggesting that the lack of calmodulin binding by CFTR is sufficient to decrease the function of PMCA4. Interestingly, dSTORM analysis of HeLa cells transfected with CFTR(S768A) and PMCA4 showed no change in the localisation of the proteins relative to each other suggesting that the binding of calmodulin has no effect on the localisation of the proteins within this nanodomain (Figure 6. E.).

**Figure 6.**
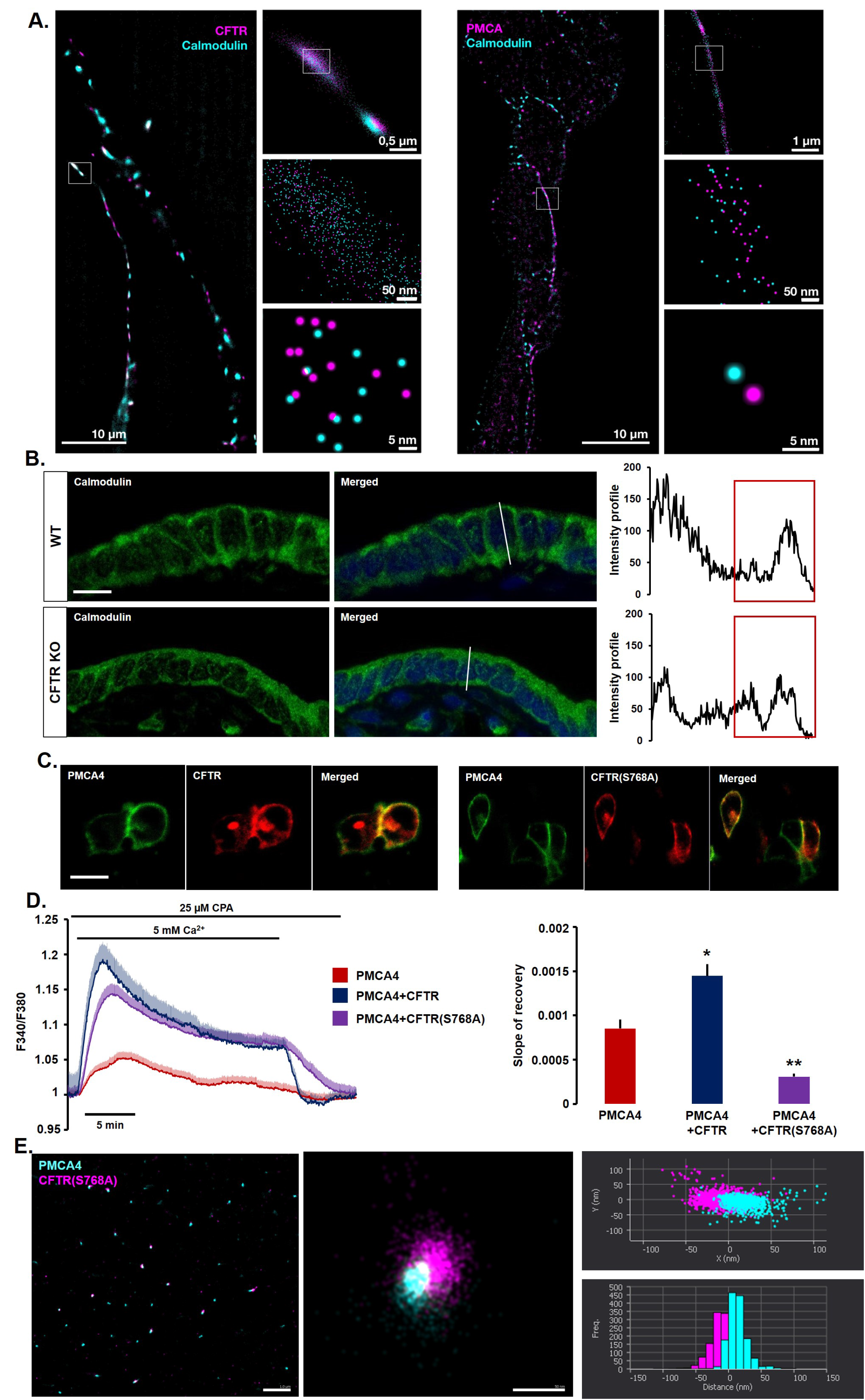
The role of CFTR-mediated calmodulin binding in the regulation of PMCA4 activity in pancreatic ductal cells. **A.** dSTORM images demonstrate the colocalization of calmodulin with CFTR and PMCA4 on cross-sections of mouse pancreatic ductal organoids. **B.** Confocal images of the intracellular distribution of calmodulin in WT and CFTR KO ductal cells. Calmodulin was strongly associated with the apical membrane in WT ductal epithelial cells, in CFTR KO cells this association was lost. Red quadrants on the line intensity profile highlight the apical region of the cells. Scale bar: 10 μm. **C.** Confocal images of WT and calmodulin binding site mutant CFTR(S768A) and PMCA4 in transfectedHEK-293 cells. Scale bar: 10 μm. **D.** Average traces and bar chart of the slope of Ca^2+^ extrusion in transfected HEK-293 cells. Cotransfection of PMCA4 and CFTR markedly increased the slope of Ca^2+^ extrusion, on the other hand CFTR(S768A) significantly impaired the activity of PMCA4. All averages were calculated from 6-10 individual experiments. *: p< 0.05 vs PMCA4, **: p< 0.05 vs PMCA4+CFTR. **E.** dSTORM images of HeLa cells transfected with CFTR(S768A) and PMCA4. The expression of the mutant CFTR had no effect on the colocalization of CFTR and PMCA4. Scale bars: 1 μm and 50 nm respectively. n= 5-7 cells were analysed for each condition in dSTORM.

### Mitochondrial function is impaired whereas apoptosis is increased in CFTR KO pancreatic ducts

The sustained intracellular Ca^2+^ elevation can impair mitochondrial function leading to apoptosis. To dissect, whether the decrease of intracellular Ca^2+^ extrusion observed in CFTR KO ductal epithelial cells is sufficient to damage mitochondria, we analysed the mitochondrial morphology and function in WT and CFTR KO ductal cells. Using transmission electron microscopy we found, that the mitochondrial volume/cell volume was not significantly different in CFTR KO ductal cells compared to WT, demonstrating that morphology of the mitochondria is not altered in CFTR KO cells (Figure 7. A.). Next, we followed the changes of mitochondrial membrane potential (Δψ_m_) during carbachol stimulation to visualize how the increased intracellular Ca^2+^ concentration alters mitochondrial function. The administration of 100 μM carbachol resulted in a permanent decrease in the Δψm in the CFTR KO, but not in WT ductal cells suggesting that the sustained elevation of intracellular Ca^2+^ leads to an impaired mitochondrial function (Figure 7. B.). ATP generated during mitochondrial oxidative phosphorylation provides fuel for ATPases such as PMCA. To rule out that the general decrease in ATP production cause the impaired function of PMCA we used oligomycin to inhibit the F1F0 ATP synthase in WT ducts. In these experiments we were unable to observe any difference in Ca^2+^ extrusion (Supplementary Figure 4.) illustrating that the impaired mitochondrial ATP production has no effect on PMCA function. Sustained Ca^2+^ elevation and disturbed mitochondrial function may lead to apoptosis. To test this, we analysed the intracellular distribution of cytochrome c, as the release of cytochrome c from the mitochondria is a hallmark of apoptosis. Indeed, the cytoplasmic intensity of cytochrome c significantly increased in the CFTR KO pancreatic ductal cell (Figure 7. C.). In addition, we also found that the cytoplasmic intensity of the initiator caspase 9 was also increased in CFTR KO cell further confirming the increased rate of apoptosis (Figure 7. D.).

**Figure 7.**
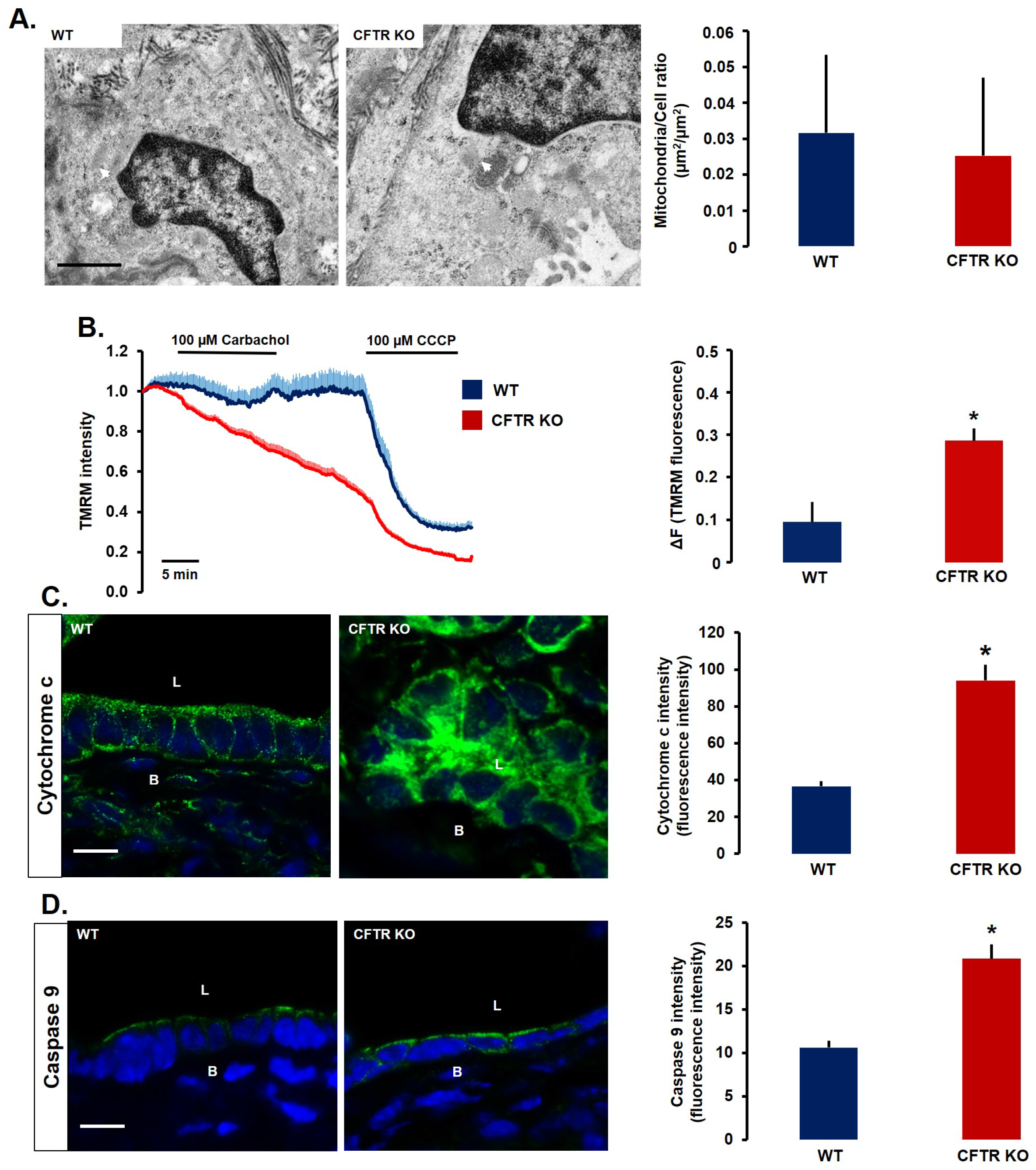
Mitochondrial dysfunction and increased expression of apoptotic proteins in CFTR KO pancreatic ductal cells. **A.** Representative transmission electron microscopy images of WT and CFTR KO mouse ductal cells (arrowheads highlight the mitochondria). The mitochondrial volume/cell volume was not changed in CFTR KO ductal cell compared to WT. Scale bar: 1 μm. **B.** Average traces of changes in mitochondrial membrane potential (Δψ_m_) and bar charts of maximal fluorescent intensity changes in response to 100 μM carbachol. Intracellular Ca^2+^ concentration increase resulted in a permanent decrease in the Δψm in the CFTR KO, but not in WT ductal cells. All averages were calculated from 6-10 individual experiments. *: p< 0.05 vs WT. **C.** Intracellular distribution of cytochrome c and **D.** caspase 9 staining in WT and CFTR KO ductal cells. In WT ductal cells cytochrome c showed a granular pattern, whereas in CFTR KO cells this was changed to a cytosolic staining. The intensity of caspase 9 staining was increased in CFTR KO cells. *: p< 0.05 vs WT. Scale bars: 10 μm. B: basolateral side; L: lumen.

## Discussion

Although the life expectancy of CF patients remained unacceptably low, with novel therapies such as potentiator-corrector combinations, the GI manifestations of the disease came to the spotlight. CF patients are at an increased risk to develop AP leading to chronic damage of the exocrine pancreas. Research studies so far focused on the consequences of impaired ion secretion by CFTR^22^, however the decreased expression of the channel might induce subcellular changes that can enhance the damage of the pancreas.

Traditionally CFTR is considered as a cAMP-activated Cl^-^ channel and the regulation of CFTR activity by intracellular Ca^2+^ levels, or the possible interactions of the channel with components of the Ca^2+^ signalling that shape subcellular signalling events is not well understood^23,24^. In our study, we demonstrated that the Ca^2+^ extrusion is significantly impaired in CFTR KO ductal cells due to the diminished function of PMCA. Notably, the Ca^2+^ extrusion was not altered in the pancreatic acinar cells isolated from the same CFTR KO mouse suggesting that the decreased Ca^2+^ extrusion is caused by the lack of CFTR function, or expression in ductal cells rather than by an unspecific injury of the exocrine pancreas. The lack of CFTR function might influence the membrane potential affecting Ca^2+^ extrusion. To test this possibility, we pharmacologically inhibited CFTR function, which had no effect on the Ca^2+^ extrusion indicating that the observed changes can be attributed to the lack of CFTR expression. We showed that NCX has no contribution to the Ca^2+^ extrusion in ductal cells proving that impaired Ca^2+^ extrusion is solely due to the dysfunction of PMCA. To rule out that this effect is species or model specific, we confirmed the PMCA dysfunction in multiple independent model systems that included siCFTR-treated pancreatic ductal fragments, heterologous expression of CFTR and PMCA in HEK-293 cells, and CFPAC-1 cells transfected with human CFTR gene. Most importantly, we also demonstrated the impaired function of PMCA in iPSC-derived human pancreatic organoids from CF patients. Previously, in cultured bronchial epithelial cells that express F508del CFTR, altered intracellular Ca^2+^ signalling was described due to elevated IP3R-dependent Ca^2+^ release, increased activity of SERCA and decreased function of PMCA^16,17^ In addition, mitochondrial Ca^2+^ uptake was enhanced in CF cells compared to controls^25^. Correction of the CFTR expression with VX809 in this model seemed to restore the alterations of the intracellular Ca^2+^ signalling^26^. Besides increased intracellular Ca^2+^ release and impaired Ca^2+^ clearance, increased activity of the extracellular Ca^2+^ influx was also described in CF. Antigny et al. demonstrated that transient receptor potential canonical 6 (TRPC6) channel-dependent extracellular Ca^2+^ influx is increased in CF airway cells^27^. Moreover, Balghi et al. showed that Orai1-mediated extracellular Ca^2+^ influx is also increased in CF airway cells leading to increased secretion of the proinflammatory cytokine IL-8^28^. Previously, our group demonstrated that sustained intracellular Ca^2+^ overload in pancreatic ductal cells is a hallmark of AP, which is a common feature regardless of the aetiology of the disease^29,30^. Common toxins, such as bile acids, or non-oxidative ethanol metabolites can trigger the release of ER Ca^2+^ stores and activate extracellular Ca^2+^ influx leading to impaired fluid and HCO_3_^-^ secretion. The sustained intracellular Ca^2+^ overload also triggers mitochondrial damage with consequent ATP depletion and cell damage^31^ that further impair the function of the ATP-dependent Ca^2+^ extrusion. However, prevention of the toxic Ca^2+^ signal generation maintains the ductal function and decreases the severity of AP. These observations suggest that protection of the ductal cell in CF from the prolonged intracellular Ca^2+^ elevation might improve exocrine pancreatic function in CF^1^.

In our experiments, we demonstrated that PMCA4 colocalizes with CFTR at the apical membrane of the ductal cells in multiple models. Moreover, using super-resolution dSTORM technique we also highlighted that CFTR and PMCA4 are within 20 nm distance suggesting a physical interaction of the two proteins. Previously, Philippe et al. suggested based on co-immunoprecipitation experiments that CFTR interacts with SERCA and PMCA in airway epithelial cells, however the nature of this interaction was not revealed^17^ Recently, Bozoky et al. showed that calmodulin binds directly to the R domain of CFTR increasing the open probability of the channel^21^, which provided a novel mechanism for the regulation of CFTR activity by intracellular Ca^2+^. The authors also demonstrated that calmodulin binds to CFTR in an alternative binding conformation allowing the two lobes of calmodulin to bind to two separate sequences independently. This binding conformation may allow CFTR to recruit calmodulin and determine the activity of other calmodulin regulated proteins such as PMCA isoforms in a macromolecular complex at the apical plasma membrane. Ca^2+^-calmodulin binds to the calmodulin binding domain of PMCA and competitively antagonizes the autoinhibitory domain leading to PMCA activation^32^. In pancreatic ductal epithelia calmodulin was associated with the apical membrane and strongly colocalized with CFTR and with PMCA4 in WT ductal cells. This apical localization was lost in CFTR KO ductal cells suggesting that CFTR is necessary to recruit calmodulin. To prove that calmodulin binding of CFTR is required for PMCA activation, we overexpressed calmodulin binding site mutant CFTR(S768A) and PMCA4 in HEK-293 cells and showed that the Ca^2+^ extrusion is significantly impaired in the presence of the mutant CFTR. The R domain connects CFTR to several other plasma membrane transporter and cytosolic structural proteins. Among others, CFTR interacts with the electrogenic SLC26 (presumably A6 and A3) Cl^-^/HCO_3_^-^ exchangers^9,33^, which promotes the secretion of HCO3^-^ to the pancreatic ductal lumen. Among structural proteins, the interaction of CFTR with the Na^+^/H^+^ exchanger regulatory factor-1 (a scaffolding protein that directs proteins to the apical plasma membrane) was previously described^34^, which can also bind the PDZ binding domain of PMCA^35^. This might explain why we were not able to disrupt the colocalization of CFTR and PMCA4 when calmodulin binding site of CFTR was mutated.

During the pathogenesis of AP it is well-established that sustained intracellular Ca^2+^ elevation induces the opening of the mitochondrial permeability transition pore, dissipating ΔΨ_m_ with a consequent drop of ATP synthesis and increasing the permeability of the inner mitochondrial membrane that will result in mitochondrial swelling and cell necrosis^36-40^. In our experiments, we found no evidence for morphological damage of the mitochondria, however upon challenging the ductal fragments from CFTR KO mice we detected a remarkable drop of ΔΨm. In addition, cytosolic staining for cytochrome c and caspase 9 were increased in CFTR KO ductal cells, suggesting an increase in the activity of apoptosis. These results suggest that the sustained Ca^2+^ elevation due to the PMCA dysfunction can contribute to the pancreatic damage in CF.

Taken together, in this study we analysed for the first time the subcellular changes in the pancreatic ductal epithelia in CF and provided evidence that the intracellular Ca^2+^ homeostasis is disturbed by the decreased expression of CFTR channel in CF ductal cells. More specifically, using multiple independent model systems, we described the decreased activity of PMCA4, which colocalizes and physically interacts with CFTR on the apical membrane of the ductal cells. The regulation of PMCA4 activity by CFTR involves recruitment of calmodulin to the apical membrane by CFTR. Even more importantly, using human iPSC-derived pancreatic organoids from healthy subjects and CF patients we observed the same decrease in PMCA4 activity highlighting the human relevance of our results. In addition, we showed that elevation of the intracellular Ca^2+^ level in CFTR KO ductal cells impaired the mitochondrial function accompanied by cytochrome c release and increased expression of caspase 9 indictaing increased activity of apoptosis. Based on these results, the prevention of sustained intracellular Ca^2+^ overload may improve the exocrine pancreatic function in CF and may be a potential therapy to prevent CF-related AP episodes and the development of pancreatic diabetes.

## Materials and methods

All materials with catalogue numbers and manufacturers used in the study are listed in Supplementary Table 1.

### Cell lines and animals

HEK-293 and HeLa cells (ATCC; Cat. No.: ATCC-CRL-3249 and ATCC-CCL-2, respectively) were grown in DMEM/F12 containing 10% fetal bovine serum (FBS), 1 % Kanamycin, 1 % Penicillin-Streptomycin and 1% GlutaMax supplement^41^. CFPAC-1 cells were a generous gift of Michael Gray (School of Biomedical, Nutritional and Sport Sciences; Newcastle University) and were grown in Iscove’s modified Dulbecco’s medium supplemented as described above. Transfection of CFPAC-1 cells with recombinant Sendai virus containing the human CFTR gene was performed as described previously^19^. CFTR KO mice were originally generated by Ratcliff et al. and were kind gift of Professor Ursula Seidler^42^. The mice used in this study were 8-12 weeks old and weighted 20-25 grams, the gender ratio was 1:1 for all groups. Experiments were carried out with adherence to the NIH guidelines and the EU directive 2010/63/EU for the protection of animals used for scientific purposes. The study was authorized by the National Scientific Ethical Committee on Animal Experimentation under licence number (XXI./2523/2018.).

### Isolation of pancreatic ductal fragments and acinar cells

Pancreatic ductal fragments were isolated as described earlier^14,43^. Briefly after terminal anaesthesia with pentobarbital the pancreas was surgically removed from the animals and placed into ice-cold DMEM/F12^14^. The pancreas was injected with 100 U/ml collagenase, 0.1 mg/ml trypsin inhibitor, 1 mg/ml bovine serum albumin in DMEM/F12 and placed into a shaking water bath at 37°C for 30 min. Small intra-/interlobular ducts were identified and isolated under stereomicroscope. For pancreatic acinar cell isolation, the tissue was injected with 200 units/ml type 4 collagenase in standard HEPES and placed into a shaking water bath at 37°C for 30 min as described previously^44^ The tube was vigorously shaken every 5 min. The supernatant was centrifuged for 2 min at 800 rpm and the pellet was resuspended in Media 199 with 0.1% BSA.

### Mouse and human pancreatic ductal organoid culture

Mouse pancreatic ductal organoids were generated as previously described with modifications^45,46^. Human pancreatic tissue samples were collected from transplantation donors (Ethical approval No.: 37/2017-SZTE). Briefly, mouse and human tissues were minced into small fragments and incubated for 1 h at 37°C in digestion solution. After this the tissue was washed and resuspended in TrypLE™ Express Enzyme 1x for 15 min at 37°C followed by resuspension in Matrigel Basement Membrane Matrix. Organoids were used for experiments between passage number 1-5. The composition of different media is listed in Supplementary table 2–5.

### Generation of human CF-specific induced PSCs

Reprogramming of keratinocytes isolated from plucked hair of a CF patient or a healthy donor was performed to generate CF-specific and control induced pluripotent stem cells as previously described^20^. The pluripotent state of the established iPS cell lines was validated by immunostaining of essential pluripotency markers including OCT4, NANOG and SSEA4, and by transcriptional profiling. DNA sequencing confirmed the patient-specific CFTR gene alteration in the respective iPSCs. The CF patient harboured the compound heterozygous mutations p.F508del and p.L1258Ffs*7.

### Constructs and transfection

HEK-293 cells were transfected with plasmids coding EGFP-hPMCA4b, mCherry-CFTR-3xHA and mCherry-CFTR-3xHA(S768A). Transfection was carried out using Lipofectamine2000 as described earlier^41^. EGFP-hPMCA4b was ordered from Addgene (Plasmid #: 47589). CFTR-3xHA was a generous gift from Gergely Lukács (McGill University, Montreal, Canada)^14^, which was then cloned into a mCherry vector. Calmodulin binding site on human CFTR was disrupted by introducing a point mutation at the position 768 using Q5 Site-Directed Mutagenesis Kit resulting in a switch from serine to alanine (Fwd: AAGGAGGCAG**<u>G</u>**CTGTCCTGA, Rev: CGTGCCTGAAGCGTGG)^21^. Anti-CFTR siRNA and transfection control were used for *cftr* silencing. The ducts were transfected with 50 nM siCFTR or siGLO Green transfection indicator in Opti-MEM then left to incubate for 12 h and were used for immunofluorescent staining or *in vitro* Ca^2+^ measurements.

### Fluorescent microscopy

Intracellular Ca^2+^ concentration ([Ca^2+^]_i_) or intracellular pH were measured as described earlier^46^ by loading the cells with Fura-2-AM, or with BCECF-AM, respectively. Ducts or acini were attached to a poly-L-lysine-coated coverslips and mounted on an Olympus IX71 fluorescent microscope equipped with an MT-20 illumination system. Filter sets for BCECF and Fura-2 were described previously^46^. The signal was captured by a Hamamatsu ORCA-ER CCD camera trough a 20X oil immersion objective (Olympus; NA: 0.8) with a temporal resolution of 1 s. Changes of ΔΨ_m_ was followed by 100 nM tertamethylrhodamine-methyl ester (TMRM) using a Zeiss LSM880 confocal microscope (excitation: 543 nm; emission: 560-650nm). The fluorescence signal was normalized and was expressed as relative fluorescence.

### Gene expression analysis

Total RNA was isolated from whole pancreatic tissue or from isolated ductal fragments^46^. The isolated RNA was reverse transcribed and amplicons were detected by ABI PRISM 7000 using SyberGreen. Beta-2 microglobulin (B2M) and Proteasome Subunit Beta 6 (PSMB6) were selected for reference genes. Relative gene expression analysis was performed by ΔΔCq technique.

### Immunfluorescent labelling and Doulink Proximity Ligation Assay

Isolated pancreatic ducts, or organoids were frozen in Shandon Cryomatrix and were sectioned and stained as previously described^46^. Cell lines were grown on cover glass and fixed without sectioning. Sections were fixed in 4% paraformaldehyde in phosphate buffered saline for 15 min then washed in 1x Tris-buffered saline (TBS) for 3 x 5 minutes. Antigen retrieval (AGR) was performed, and sections were blocked with 0.1% goat serum and 10% BSA in TBS for 1 h. Incubation with primary antibodies was performed overnight at 4°C then secondary antibody was added for 2h at room temperature. Applied antibodies are listed in Supplementary Table 6–7. For Proximity Ligation Assay (PLA) guinea pig pancreatic ducts were isolated, cryosectioned and fixed as described above. AGR was performed as described above. Duolink^®^ assay [Sigma Aldrich Cat. No.: DUO92014, DUO82049, DUO92001, DUO92005] was performed on a humidified chamber according to the protocol provided by the manufacturer. Nuclear staining was performed with 1 μg/ml Hoechst33342 for 15 min and sections were mounted with Flouromount. Images were captured with a Zeiss LSM880 confocal microscope using a 40X oil immersion objective (Zeiss, NA: 1.4).

### Direct Stochastic Optical Reconstruction Microscopy (dSTORM)

HeLa cells grown on cover glass were co-transfected with EGFP-hPMCA4b, mCherry-CFTR-3xHA and mCherry-CFTR-3xHA(S768A) plasmids. Primary human pancreatic ductal cells were isolated from transplantation donors and were generated for dSTORM by digesting and plating human pancreatic ductal organoids on cover glass. HeLa cells were fixed with 4% PFA for 10 min and AGR was performed by 0.01% Triton-X-100 for 10 min. Applied antibodies are listed in Supplementary Table 7. Adherent primary cell and sectioned organoids were fixed and retrieved by methanol for 10 min at −20 °C. Cover glasses were placed in blinking buffer and dSTORM images were captured by Nanoimager S (Oxford Nanoimaging ONI Ltd.).

### Transmission Electron Microscopy

The pancreatic tissue was fixed in 2% glutaraldehyde supplemented with 2.25% dextran in phosphate buffered saline overnight at 4°C^31^. The tissue was embedded into Embed812 and 70 nm thick ultrathin sections were cut with a Leica ultramicrotome. For determination of mitochondrial morphology 18-30 cells in each sample were captured on a Jeol 1400 plus electron microscope with a magnification of 12000X. The volume fraction of the cells and mitochondria were measured by point counting, where the area per test point was 500nm×500nm for cell and 50nm×50nm for mitochondria. The results were given as the ratio of mitochondrial volume/cell volume.

### Statistical analysis

Data are expressed as means ± SE. Significant difference between groups was determined by Mann-Whitney test. *P* < 0.05 was accepted as being significant.

## Acknowledgements

The research was supported by funding from the Hungarian National Research, Development and Innovation Office (GINOP-2.3.2–15–2016–00048 to JM, PD116553 to PP), the Ministry of Human Capacities (EFOP 3.6.2-16-2017-00006 to JM), Bolyai Research Fellowship (BO/00569/17 to PP), the Hungarian Academy of Sciences (LP2017–18/2017 to JM), by the National Excellence Programme (20391-3/2018/FEKUSTRAT to JM), by the New National Excellence Program of the Ministry of Human Capacities (UNKP-18-4-SZTE-85 to PP) and EFOP 3.6.3-VEKOP-16-2017-00009 to MT and VÁ. This work was supported by Albert Szent-Györgyi Research Grant (to MJ and PP) by the Faculty of Medicine, University of Szeged. The project has received funding from the EU’s Horizon 2020 research and innovation program under grant agreement No. 739593.

## Author contributions

TM and JM designed the research project; TM, ÁV, NP, BD, BT, PP, JF, RZ, ZR, MH, AK, MG and PH contributed to acquisition, analysis and interpretation of data for the work; TM and JM drafted the work, PH and ZR revised the manuscript critically for important intellectual content. All authors approved the final version of the manuscript, agree to be accountable for all aspects of the work in ensuring that questions related to the accuracy or integrity of any part of the work are appropriately investigated and resolved. All persons designated as authors qualify for authorship, and all those who qualify for authorship are listed.

---

**Supplementary Table 1-.** materials used during experiments listed

| Name | Provider | Cat. No.: |
| --- | --- | --- |
| HEK 293 cell line | ATCC | ATCC-CRL-3249 |
| HELA cell line | ATCC | ATCC-CCL-2 |
| DMEM (5796) | Sigma Aldrich | D6421 |
| Fetal Bovine Serum | Sigma-Aldrich | F7524 |
| Kanamycin | ThermoFisher Scientific | 15160047 |
| Strep-Pen | ThermoFisher Scientific | 15140122 |
| GlutaMax | ThermoFisher Scientific | 35050061 |
| IMDM | ThermoFisher Scientific | 12440061 |
| DME-F12 | Sigma-Aldrich | D6421 |
| collagenase | Worthington | LS005273 |
| trypsin inhibitor | ThermoFisher Scientific | 17075029 |
| type 4 collagenase | Worthington | LS004186 |
| Media 199 | ThermoFisher Scientific | 11150059 |
| TrypLE™ Express | Gibco | 12605028 |
| corning matrigel | Corning | 354234 |
| FURA-2,AM | ThermoFisher Scientific | F1201 |
| Pluronic F-127 | ThermoFisher Scientific | P3000MP |
| BCECF,AM | ThermoFisher Scientific | B1170 |
| poly-L-lysine | Sigma-Aldrich | P4707-50ML |
| TMRM | ThermoFisher Scientific | T668 |
| LBP buffer | Macherey-Nagel | 740906.125 |
| NucleoSpin RNA Plus kit | Macherey-Nagel | 740984.25 |
| High Capacity cDNA Reverse Transcription Kit (ABI) | ThermoFisher Scientific | 4368814 |
| Maxima SYBR Green/ROx qPCR MasterMix | ThermoFisher Scientific | K0222 |
| Shandon Cryomatrix | ThermoFisher Scientific | 6769006 |
| Sodium Citrate | Sigma Aldrich | C8532 |
| Tween20 | Sigma Aldrich | P1379 |
| goat serum | Sigma Aldrich | G9023 |
| bovine serum albumin | Sigma Aldrich | A8022 |
| Hoechst33342 | ThermoFisher Scientific | 62249 |
| Flouromount | Sigma Aldrich | F4680 |
| Duolink® assay | Sigma Aldrich | DUO92014, DUO82049, DUO92001, DUO92005 |
| ON-TARGETplus Mouse siRNA, SMARTpool mouse CFTR | Dharmacon | L-042164-00-0005 |
| siGlo Green transfection indicator | Dharmacon | D-001630-01-05 |
| eGFP-hPMCAb | Addgene | 47589 |
| Lipofectamine 2000 | Invitrogen | 11668-019 |
| Embed 812 | EMS | 14120 |
| 15 ml centrifuge tube | Greiner BioOne | 188261 |
| 6-well plates | Greiner BioOne | 657160 |
| cover glass | VWR | ECN 631-1583 |
| PFA | Alfa Aesar | 43368 |
| PBS | Sigma Aldrich | P4417-100TAB |
| Triton-X-100 | Reanal | 32190-1-99-33 |
| cavity slides | Sigma Aldrich | BR475505-50EA |
| two-component adhesive | Picodent | 13 007 100 |
| glucose oxidase | Sigma Aldrich | G2133-50KU |
| catalase | Sigma Aldrich | C100 |
| glucose | Sigma Aldrich | G8270 |
| cysteamine hydrochloride | Sigma Aldrich | M6500 |
| methanol | Sigma Aldrich | 34860 |
| cerulein | Bachem | H-3220 |
| A Amylase Assay | Diagnosticum | 47462 |

**Supplementary Table 2–.** Splitting media

| Component | Manufacturer/Cat.No. | Final cc/volume |
| --- | --- | --- |
| Advanced DMEM/F-12 | Gibco, Catalog No.: 12634-010 | 500 ml |
| 1 M HEPES | Gibco, Catalog No.: 15630080 | 5 ml<br>(10 mM) |
| GlutaMax Supplement (100X) | Gibco, Catalog No.: 35050061 | 5ml<br>(1X) |
| Primocin (400X) | Invivogen, Catalog No.: ant-pm-2 | 1,25 ml<br>(1X) |

**Supplementary Table 3–.** Digestion media

| Component | Manufacturer/Cat.No. | Final cc/volume |
| --- | --- | --- |
| Splitting media | - | 20 ml |
| Collagenase IV. | Worthington, Catalog No.: LS004188 | 1250 U/ml |
| Dispase | Sigma-Aldrich, Catalog No: D4693-1G | 0,5 U/ml |
| FBS | Gibco, Catalog No: 10500064 | 0,5 ml<br>2,5% v/v |
| Trypsin inhibitor | Sigma-Aldrich, Catalog No: T9128-1G | 1mg/ml |

**Supplementary Table 4–.** Wash media

| Component | Manufacturer/Cat.No. | Final cc/volume |
| --- | --- | --- |
| Splitting media | - | - |
| FBS | Gibco, Catalog No:<br>10500064 | 2,5% v/v |
| Antibiotic-Antimycotic<br>Solution (100X) | Gibco, Catalog No.:<br>15240062 | 1X |
| Kanamycin Sulfate<br>(100X) | Gibco, Catalog No.:<br>15160047 | 1X |
| Voriconazole | TOCRIS, Catalog No:<br>3760/10 | 2 µg/ml |

**Supplementary Table 5–.**
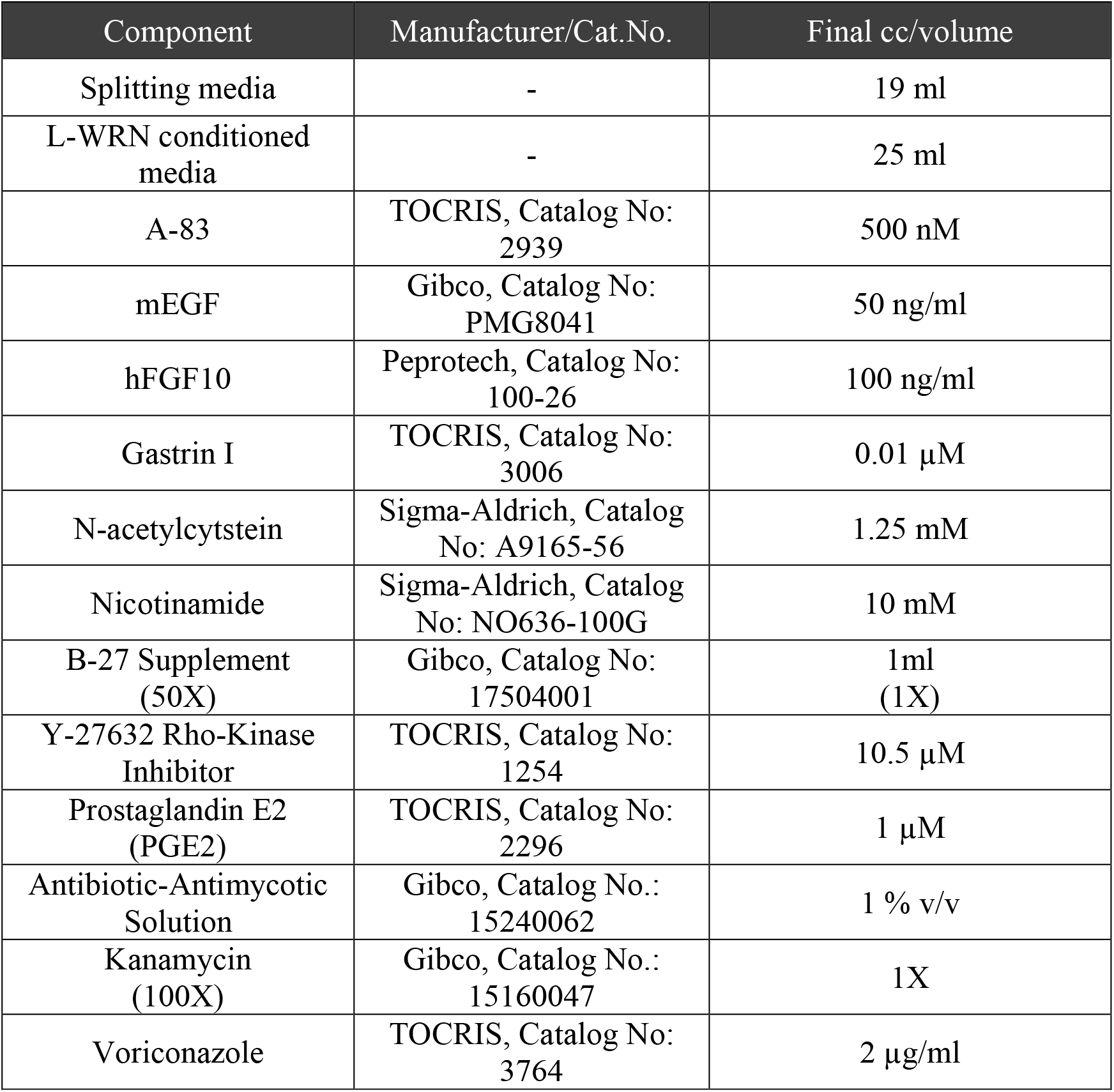
Feeding media

**Supplementary Table 6–.** Primary antibodies used during experiments:

| Name | Source | Clonality | Cat. No. | Provider |
| --- | --- | --- | --- | --- |
| **anti-CFTR** | rabbit | polyclonal | ACL-006 | Alomone Labs |
| **anti-PMCA1** | rabbit | polyclonal | ACP-005 | Alomone Labs |
| **anti- PMCA4** | rabbit | polyclonal | ACP-004 | Alomone Labs |
| **anti-CFTR** | mouse | monoclonal | ab2784 | Abcam |
| **panPMCA** | mouse | monoclonal | ab2825 | Abcam |
| **anti-cytochrome C** | mouse | monoclonal | ab13575 | Abcam |
| **anti-Calmodulin** | rabbit | monoclonal | ab45689 | Abcam |
| **anti-caspase9** | rabbit | monoclonal | ab202068 | Abcam |

**Supplementary Table 7-.** Secondary antibodies used during experiments:

| Name | Cat. No. | Provider |
| --- | --- | --- |
| Goat anti-Mouse IgG (H+L) Cross-Adsorbed Secondary Antibody, Alexa Fluor 488 | A-11001 | invitrogen |
| Goat anti-Mouse IgG (H+L) Highly Cross-Adsorbed Secondary Antibody, Alexa Fluor 594 | A-11032 | invitrogen |
| Goat anti-Rabbit IgG (H+L) Cross-Adsorbed Secondary Antibody, Alexa Fluor 568 | A-110011 | invitrogen |
| Goat anti-Rabbit IgG (H+L) Highly Cross-Adsorbed Secondary Antibody, Alexa Fluor 488 | A-11034 | invitrogen |
| Donkey anti-Mouse IgG (H+L) Highly Cross-Adsorbed Secondary Antibody, Alexa Fluor 647 | A-31571 | invitrogen |
| Goat anti-Rabbit IgG (H+L) Cross-Adsorbed Secondary Antibody, Alexa Fluor 568 | A-11011 | invitrogen |

**Supplementary Table 8-.**
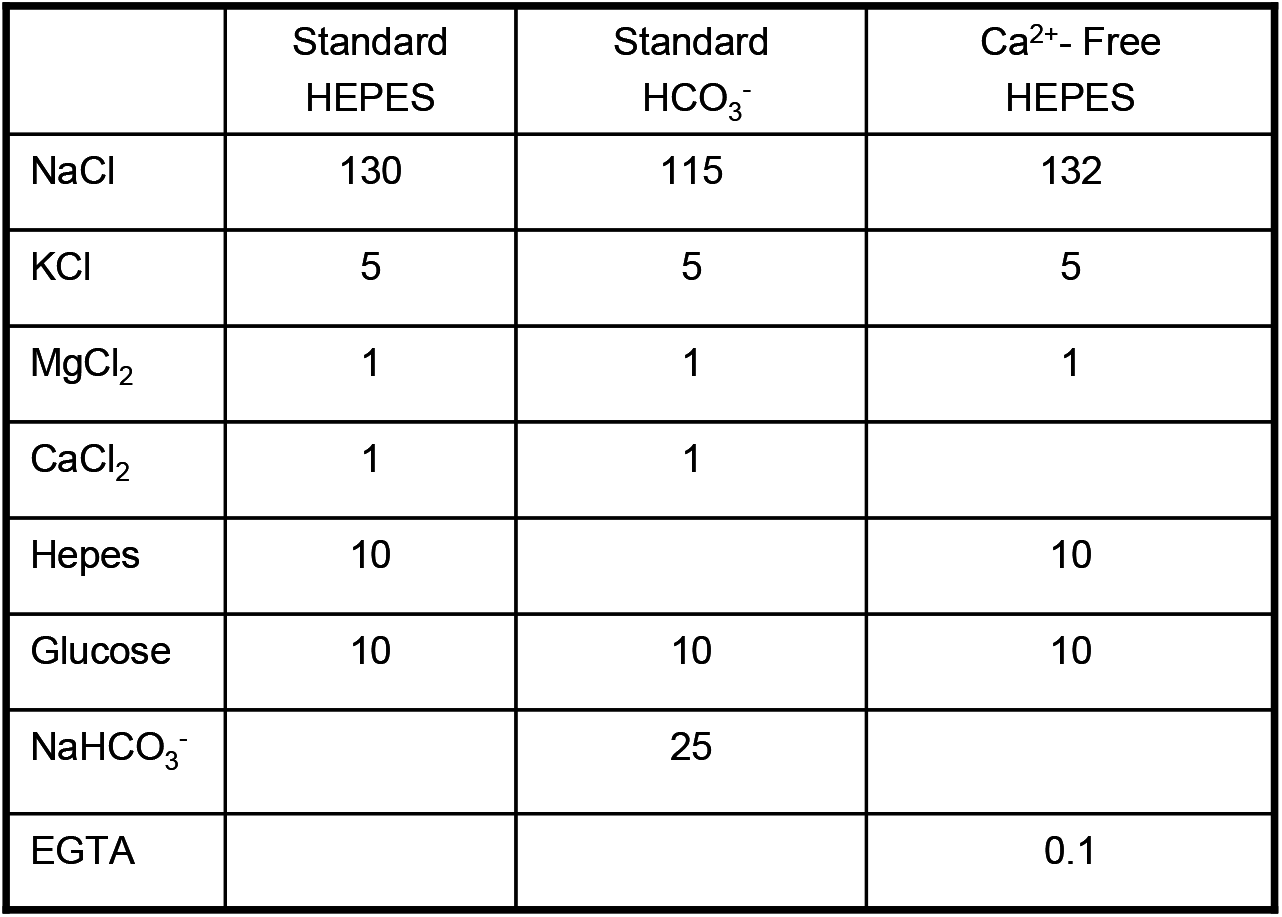
composition of solutions used during experiments

**Supplementary Figure 1.**
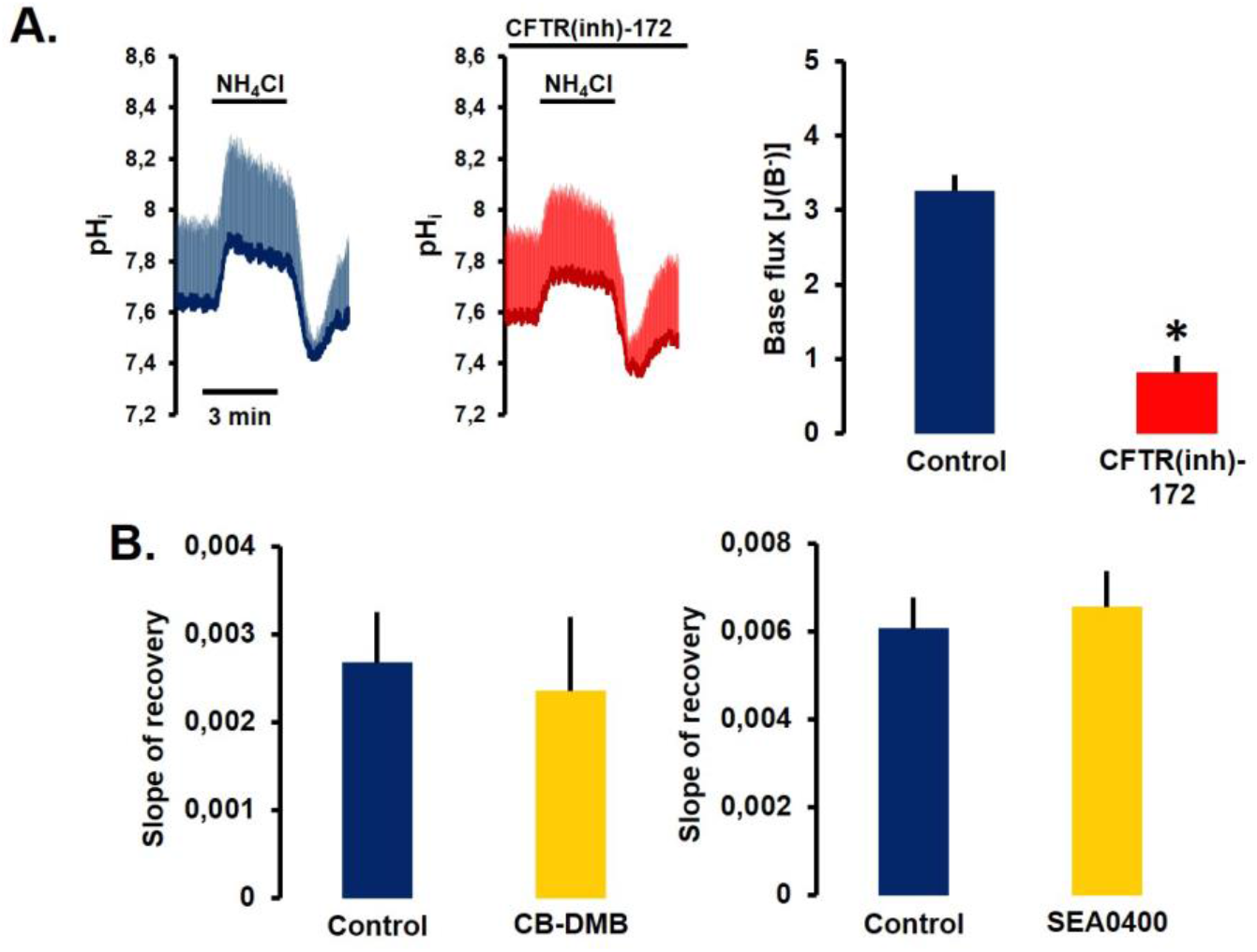
**A.** Average traces and bar charts demonstrating the effect of CFTR inhibition in isolated pancreatic ducts. The inhibition of CFTR significantly decreased the recovery from alkali load. All averages were calculated from 6-10 individual experiments. *: p< 0.05 vs Control. **B.** Bar charts of the effect of NCX inhibition in pancreatic ductal cells. CB-DMB and SEA0400 had no effect on the Ca^2+^ extrusion.

**Supplementary Figure 2.**
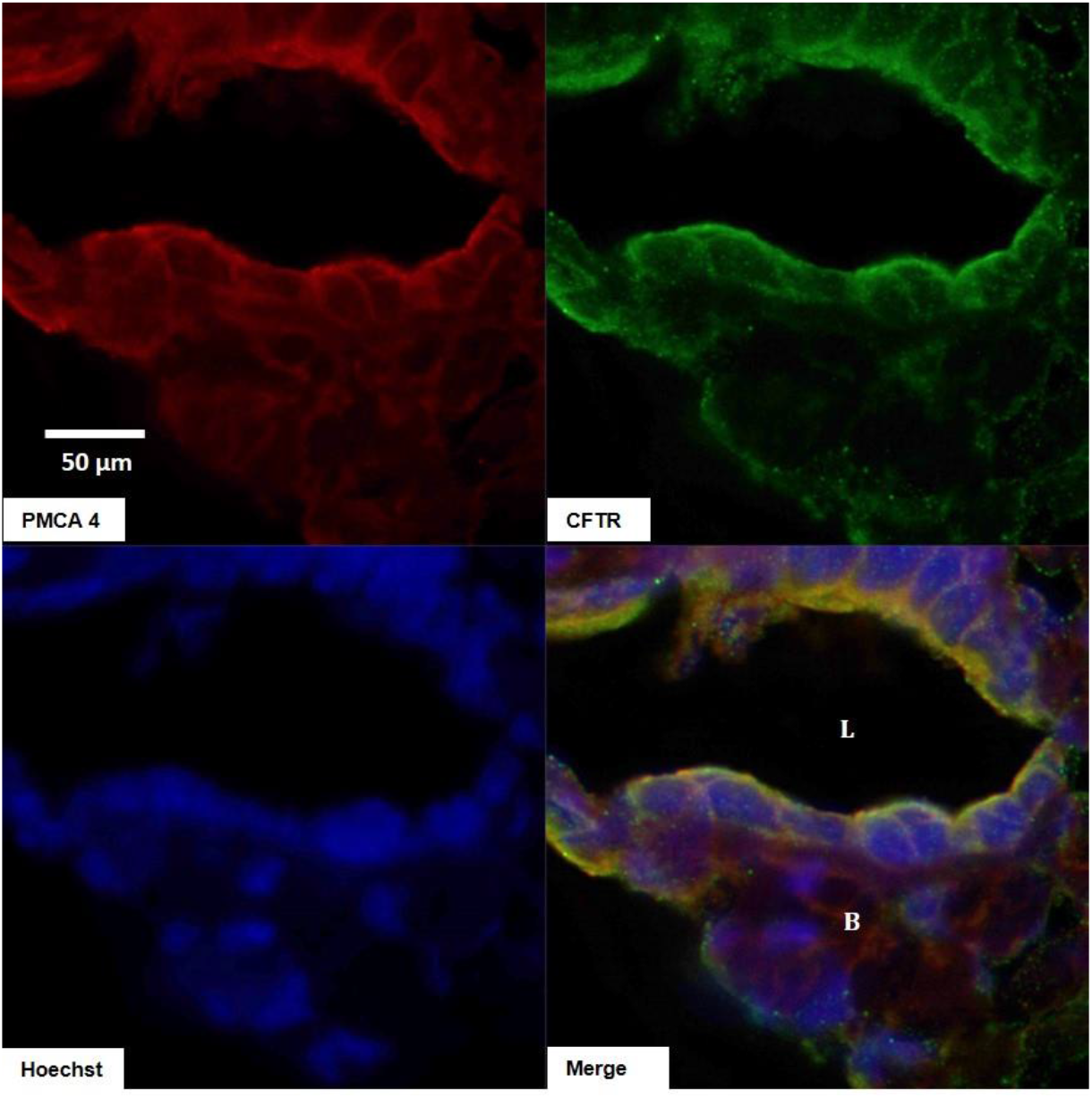
Immunofluorescent staining of CFTR and PMCA4 in guinea pig pancreatic ductal fragments.

**Supplementary Figure 3.**
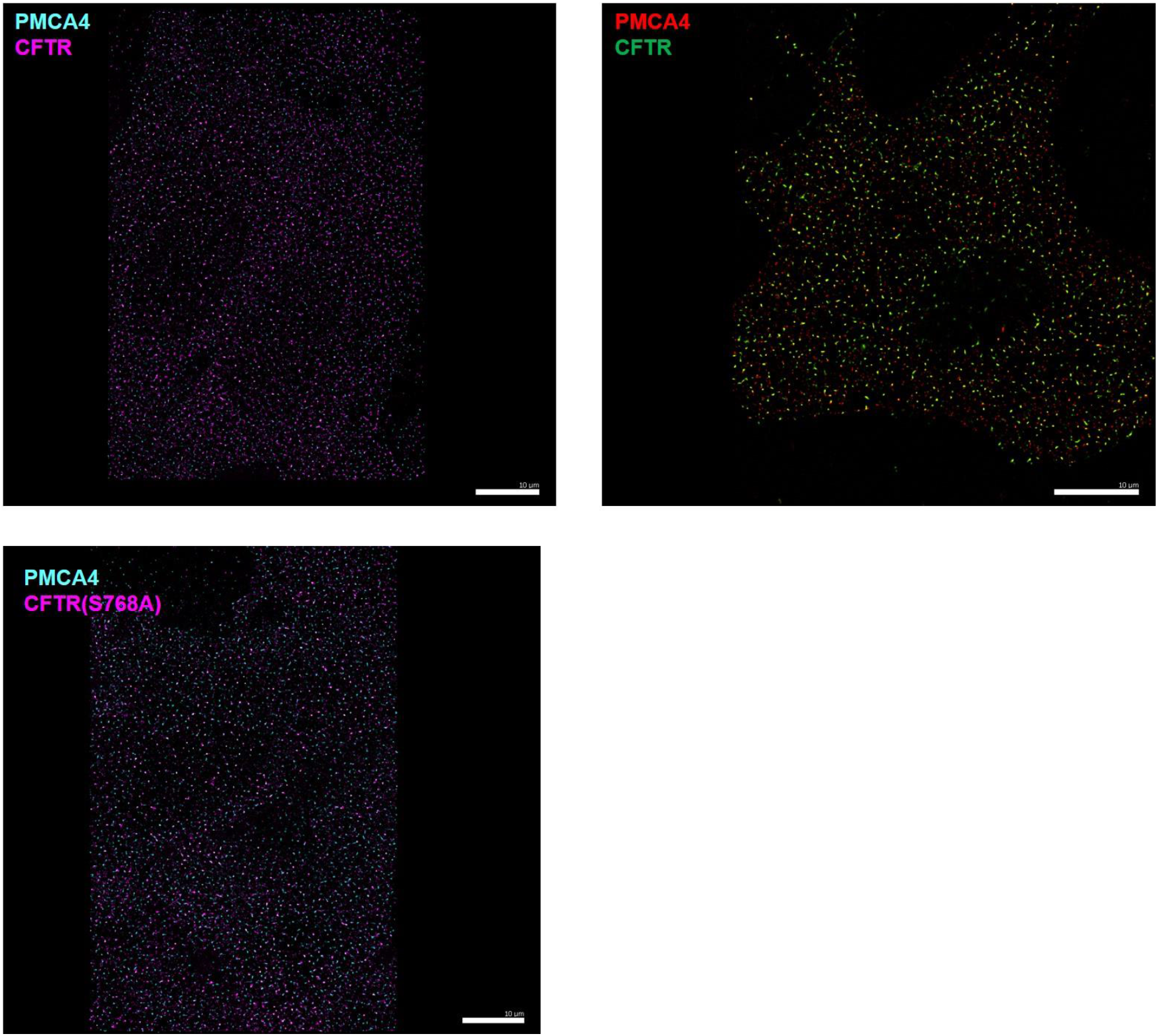
dSTORM images of PMCA4 and CFTR in transfected Hela cells (left upper and lower image) and in primary 2D human pancreatic ductal cell.

**Supplementary Figure 4.**
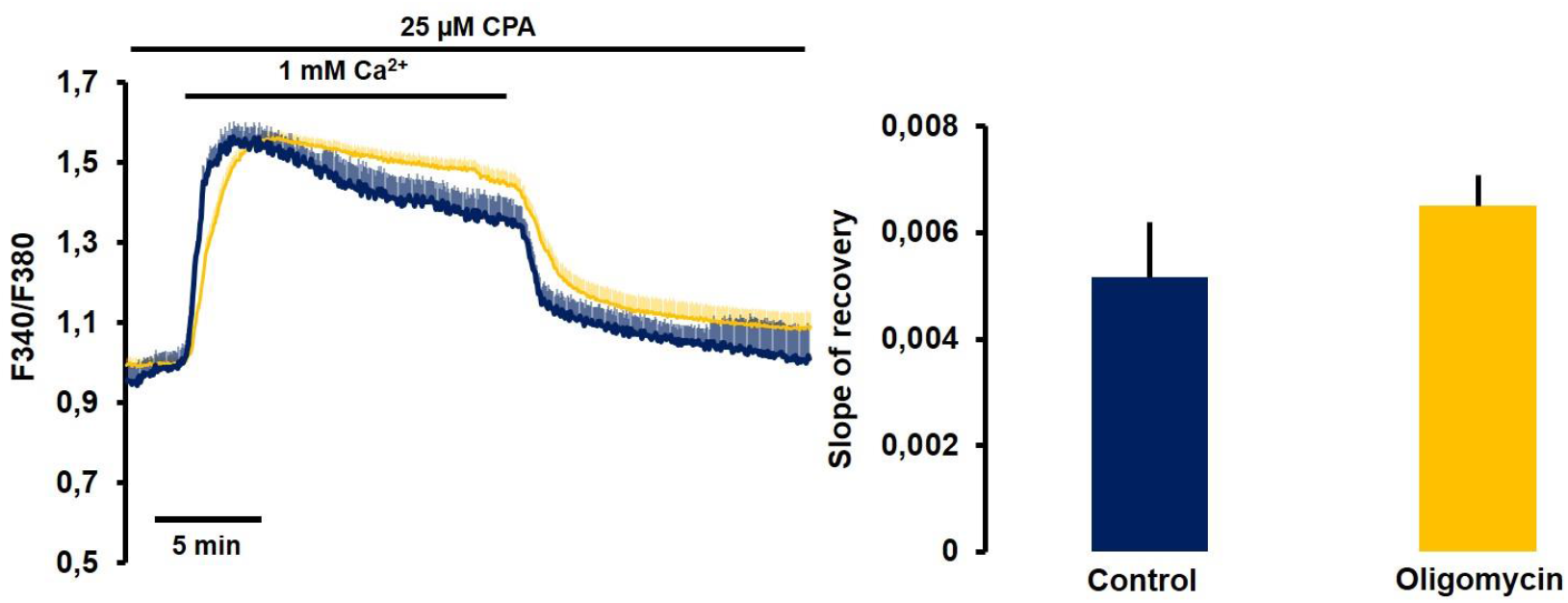
Average traces and bar charts demonstrating the effect of the inhibition of mitochondrial ATP production on the Ca^2+^ extrusion in isolated pancreatic ducts. Oligomycin had no effect on the PMCA activity. All averages were calculated from 6-10 individual experiments.

